# Maximizing genetic representation in seed collections from populations of self and cross-pollinated banana wild relatives

**DOI:** 10.1101/2021.05.25.445549

**Authors:** Simon Kallow, Bart Panis, Toan Vu Dang, Tuong Vu Dang, Janet Paofa, Arne Mertens, Rony Swennen, Steven B. Janssens

## Abstract

**Background:** Conservation of plant genetic resources, including the wild relatives of crops, plays an important and well recognised role in addressing some of the key challenges faced by humanity and the planet including ending hunger and biodiversity loss. However, the genetic diversity and representativeness of *ex situ* collections, especially that contained in seed collections, is often unknown. This limits meaningful assessments against conservation targets, impairs targeting of future collecting and limits their use.

We assessed genetic representation of seed collections compared to source populations for three wild relatives of bananas and plantains. Focal species and sampling regions were *Musa acuminata* subsp. *banksii* (Papua New Guinea), *M. balbisiana* (Viet Nam) and *M. maclayi s.l.* (Bougainville, Papua New Guinea). We sequenced 445 samples using suites of 16-20 existing and newly developed taxon-specific polymorphic microsatellite markers. Samples of each species were from five populations in a region; 15 leaf samples and 16 seed samples from one infructescence (‘bunch’) for each population.

**Results:** Allelic richness of seeds compared to populations was 51%, 81% and 93% (*M. acuminata, M. balbisiana* and *M. maclayi* respectively). Seed samples represented all common alleles in populations but omitted some rarer alleles. The number of collections required to achieve the 70% target of the Global Strategy for Plant Conservation was species dependent, relating to mating systems. *Musa acuminata* populations had low heterozygosity and diversity, indicating self-fertilization; many bunches were needed (>15) to represent regional alleles to 70%; over 90% of the alleles from a bunch are included in only two seeds. *Musa maclayi* was characteristically cross-fertilizing; only three bunches were needed to represent regional alleles; within a bunch, 16 seeds represent alleles. *Musa balbisiana,* considered cross-fertilized, had low genetic diversity; seeds of four bunches are needed to represent regional alleles; only two seeds represent alleles in a bunch.

**Conclusions:** We demonstrate empirical measurement of representation of genetic material in seeds collections in *ex situ* conservation towards conservation targets. Species mating systems profoundly affected genetic representation in seed collections and therefore should be a primary consideration to maximize genetic representation. Results are applicable to sampling strategies for other wild species.

## Background

Conservation of crop wild relatives (CWRs), wild plant species related to crops, is increasingly recognized as a vital component of both sustainable development for food security (Target 2.5 of the Sustainable Development Goals) (1) and biodiversity conservation (Target 9 of the Global Strategy for Plant Conservation) (2, 3). Importantly, this should include targeting conservation at the intraspecific level (4), essential for the functioning and flourishing of species, ecosystems (5) and crop breeding (6). Associated with policy recognition is the need for assessments against indicators or targets. However, assessment of conservation at the genetic level is often lacking and poorly understood (4, 7).

Conservation of CWRs should complementarily include both *in situ* and *ex situ* approaches (8). *Ex situ* seed conservation can maintain numerous genotypes with minimal input (9). However, knowledge of the genetic representativeness in *ex situ* seed collections, the proportion of alleles of wild populations also present in *ex situ* collections, has only been studied for a very small number of species. In fact, a recent meta-analysis of *ex situ* and *in situ* genetic comparisons only found six studies to include from seed bank collections (10). Only two of these were of wild rather than cultivated species: a Mediterranean aquatic (11), and temperate dioecious European tree species (12). There is clearly, therefore, an evidence gap for reporting on the genetic representation of species in *ex situ* seed collections.

Presently, for seed collectors to maximise genetic capture in collections, sampling guidance is often broad, encompassing all species, or inferred from taxonomically or ecologically related species (13–15). The general nature of such protocols does not always account for several key factors that shape genetics of populations and seeds, such as the existing genetic diversity of populations, the spatial distribution of plants in the environment (16–18), and species’ reproductive systems (14). It is therefore important to increase evidence-based sampling strategies to inform targeted future seed collections. Such evidence also provides valuable ecological information for *in situ* conservation and increases the value of seed collections, as it improves the selection and targeting of seed samples in breeding or phenotyping experiments.

Seed conservation of banana CWRs (*Musa* L.) is a case in point. Bananas, together with related plantains (both are *Musa*), are the most important fruit and among the most important crops in the world (19). Global production is estimated to be 116 million tonnes annually, worth $31 billion (average of 2017-19) (19). Worryingly, several biotic threats, such as by Fusarium Wilt Tropical Race 4 and Banana Bunchy Top Virus, threaten banana production. Their small genepool makes them, and the many millions of people who rely on them, particularly vulnerable (20, 21). There are around 80 taxa (hereafter referred to as ‘species’) in the genus *Musa* (22). They are tall herbaceous monocarpic monocotyledons native to tropical and subtropical Asia and the western Pacific. Most cultivated bananas and plantains derive from two species: *M. acuminata* subsp. and *M. balbisiana* (23–26). The Fe’i bananas of Pacific regions are a distinct cultivated group, deriving from *M. maclayi* (27). These focal species, included in our study (Table 1), are therefore of interest to breeders (e.g. 28).

**Table 1.**
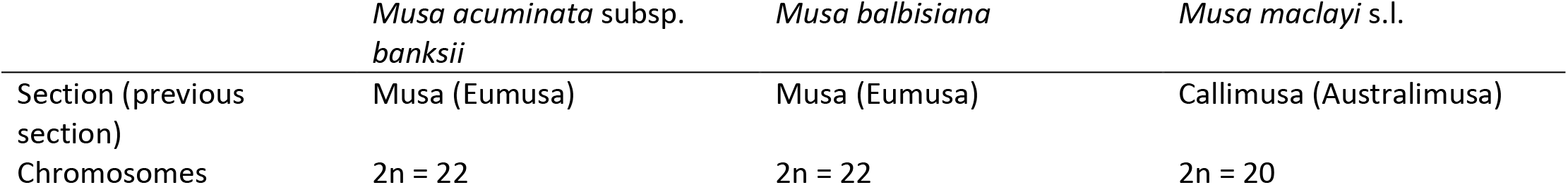
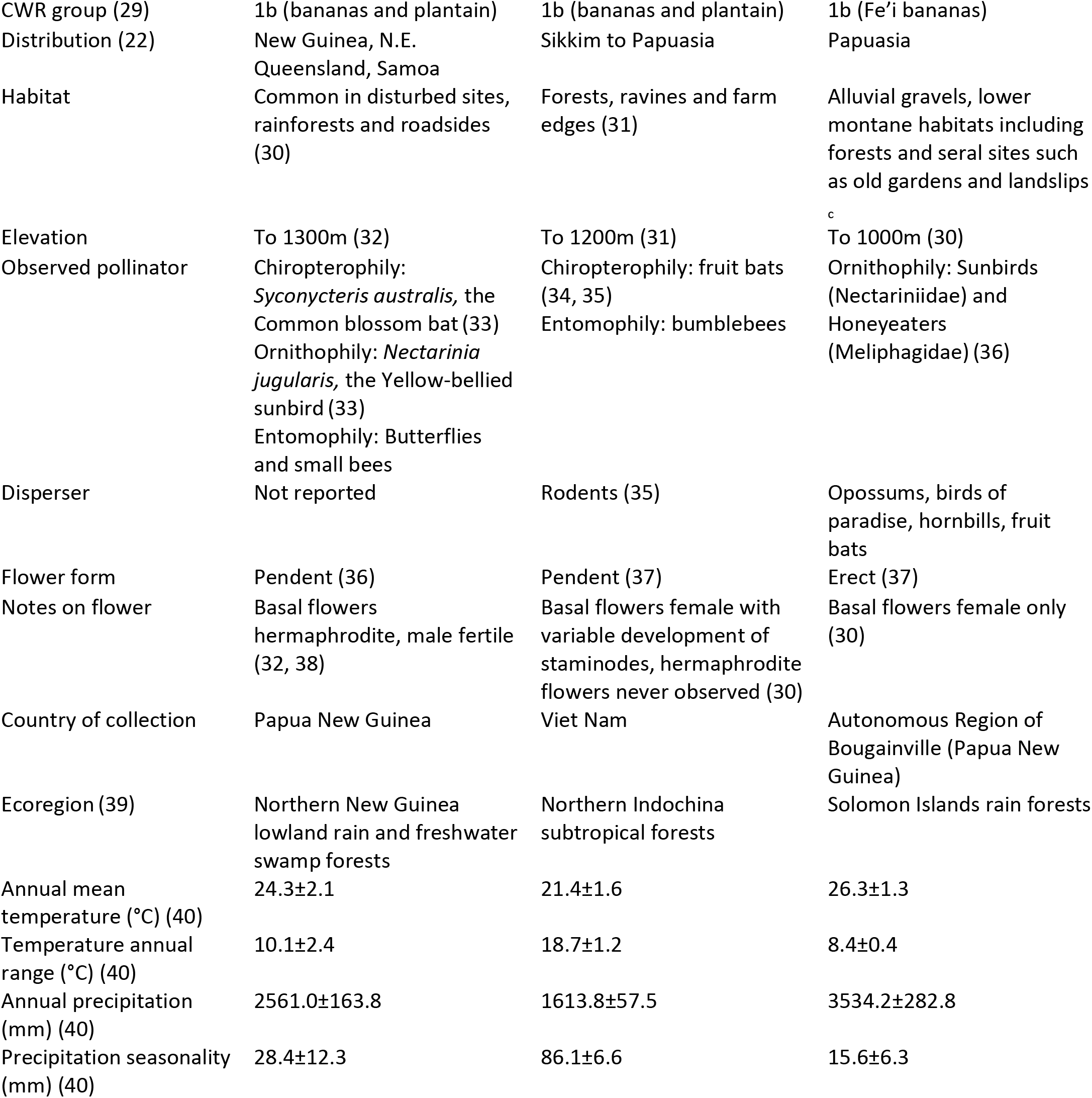
Taxonomic and ecological information on target species.

Conservation of banana CWRs is increasingly important because they are under threat. In a recent study (41), 15% of species were provisionally assessed as endangered and an additional 19% vulnerable to extinction. Furthermore, 95% of *Musa* species were assessed as insufficiently conserved *ex situ* (41).

There are only 163 genotype accessions of 35 species maintained in genebanks as living plants (42). Additionally, there are 131 seed accessions, multiple seeds collected from the same individual or population, of 10 species, stored at the Millennium Seed Bank, UK (43). Many *Musa* species are therefore not represented in genebanks at all or are represented with little or as yet unknown representativeness. An evaluation of the genetic representation of present collections will help target future conservation efforts.

The objectives of the present study are to assess and compare the genetic capture in seed collections compared to their source populations, at both regional and local scales, for three focal species; to provide guidance about how to maximize genetic capture for future seed collections; and to provide direction for seed distribution on how to provide representative seed samples.

## Results

### Representation of populations in seeds at the regional level

Allelic richness of seeds as a proportion of populations was 51%, 81% and 93% (respectively *M. acuminata, M. balbisiana, and M. maclayi*, Table 2). Allelic richness (*AR*) of populations and seeds of *M. maclayi* was much higher than the other two species. Populations had many alleles that were private (*PA*); seeds had a few *PA* - indicating that some pollination occurred by plants not present in population samples. Only two alleles in *Musa acuminata* seeds were private.

**Table 2.**
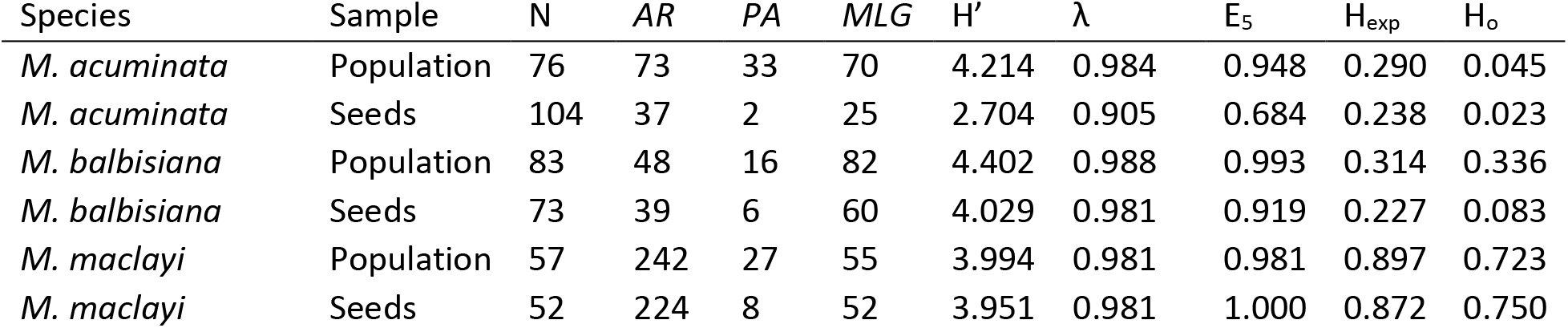
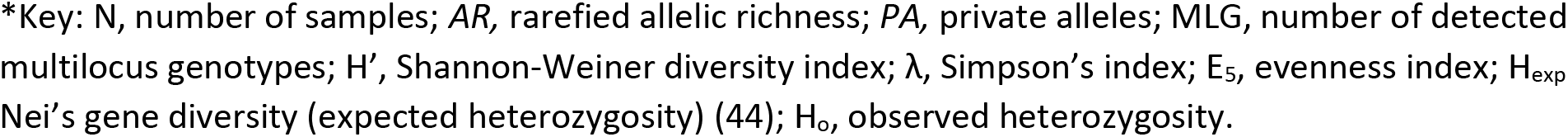
Diversity indices of populations and seeds pooled at the regional level*.

| Species | Sample | N | AR | PA | MLG | H' | $\lambda$ | E<sub>5</sub> | H<sub>exp</sub> | H<sub>o</sub> |
| --- | --- | --- | --- | --- | --- | --- | --- | --- | --- | --- |
| *M. acuminata* | Population | 76 | 73 | 33 | 70 | 4.214 | 0.984 | 0.948 | 0.290 | 0.045 |
| *M. acuminata* | Seeds | 104 | 37 | 2 | 25 | 2.704 | 0.905 | 0.684 | 0.238 | 0.023 |
| *M. balbisiana* | Population | 83 | 48 | 16 | 82 | 4.402 | 0.988 | 0.993 | 0.314 | 0.336 |
| *M. balbisiana* | Seeds | 73 | 39 | 6 | 60 | 4.029 | 0.981 | 0.919 | 0.227 | 0.083 |
| *M. maclayi* | Population | 57 | 242 | 27 | 55 | 3.994 | 0.981 | 0.981 | 0.897 | 0.723 |
| *M. maclayi* | Seeds | 52 | 224 | 8 | 52 | 3.951 | 0.981 | 1.000 | 0.872 | 0.750 |

\*Key: N, number of samples; AR, rarefied allelic richness; PA, private alleles; MLG, number of detected multilocus genotypes; H', Shannon-Weiner diversity index; $\lambda$ , Simpson's index; E<sub>s</sub>, evenness index; H<sub>exp</sub>, Nei's gene diversity (expected heterozygosity) (44); H<sub>o</sub>, observed heterozygosity.

For all species, populations were characterized by having more rare alleles than seeds; and seeds had more common alleles (Figure 2a). Seeds, therefore, captured most of the common alleles of populations, yet less so the rarer alleles. *Musa maclayi* had a notably high number of rare alleles in both populations and seeds.

**Figure 1.**
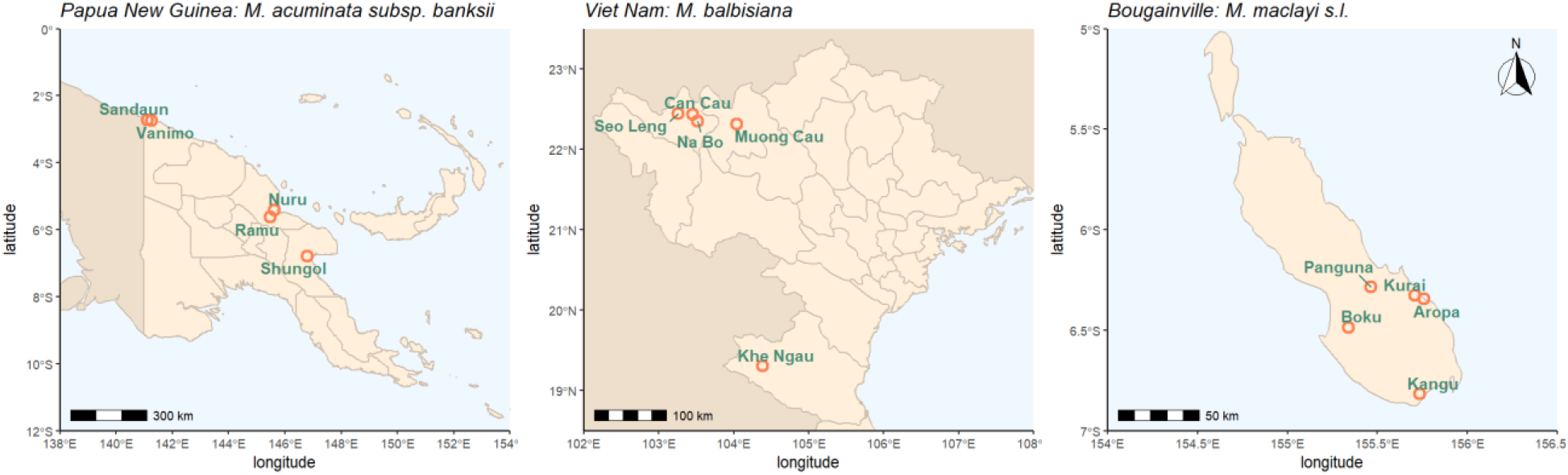
Location of populations used in this study; provinces are delineated. Map source: Natural Earth.

**Figure 2.**
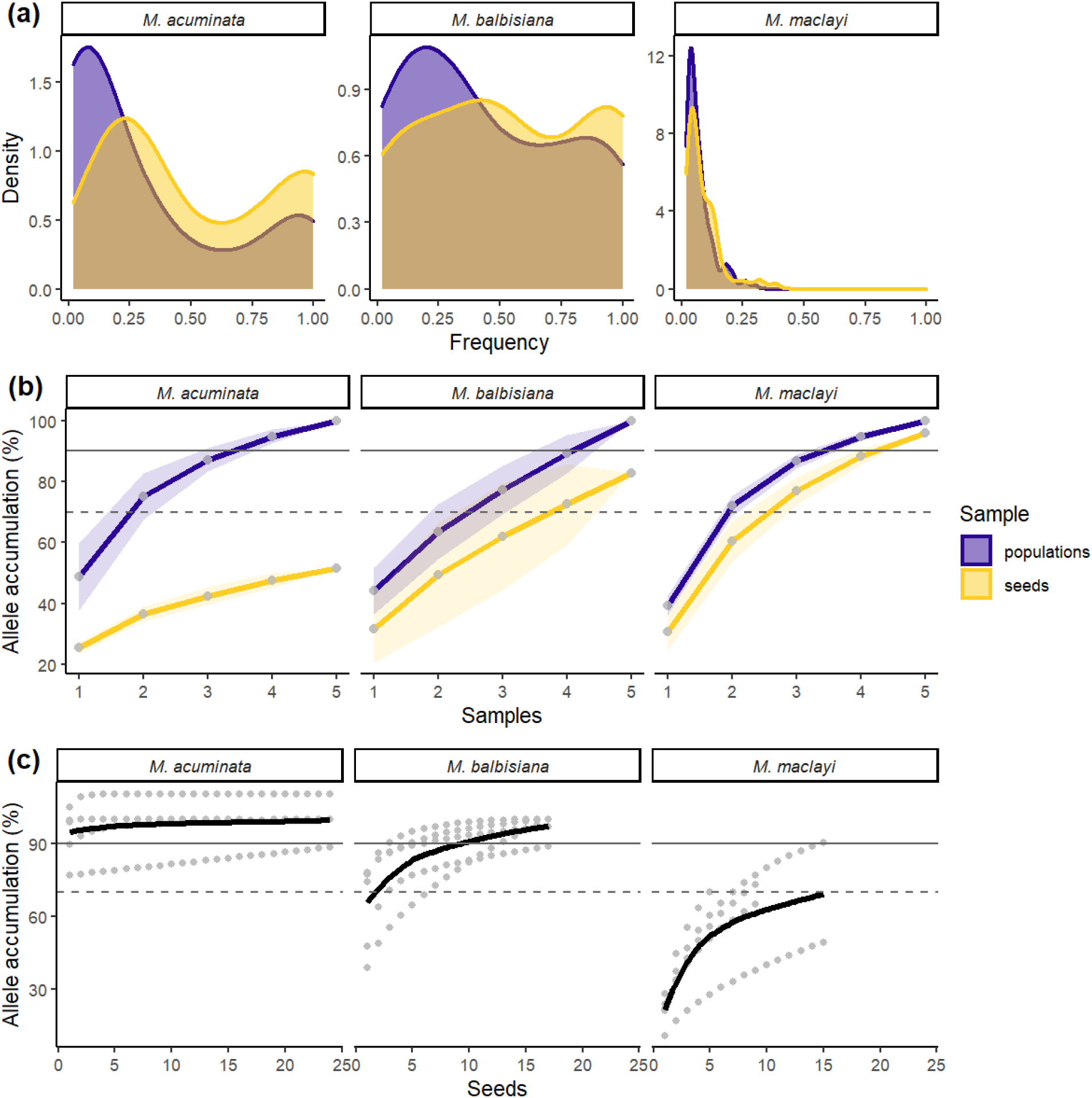
Alleles in populations and seeds of *M. acuminata, M. balbisiana* and *M. maclayi:* (a) density of allele frequencies in all populations, (b) cumulative alleles as a proportion of extrapolated total alleles in the region (shaded areas are standard deviations, (c) cumulative alleles of seeds from separate bunches, dots represent single seeds, trend line estimated using method Loess

Diversity of *M. acuminata* seeds was significantly lower than that of populations (Shannon-Weiner diversity index (H’): t=4.595, df=4.310, p=0.008; Simpson’s index (λ): t=3.163, df=4.018, p=0.034). *Musa balbisiana* seed diversity was also significantly lower than populations (H’: t=2.890, df=6.192, p=0.0267; λ: t=3.291, df=7.908, p=0.0112). However, diversity of seeds and populations of *M. maclayi* was not significantly different (H’: t=0.716, df=5.566, p=0.503; λ: t=0.817, df=5.216, p=0.450). Observed heterozygosity (H_o_) was also significantly lower in seeds compared to populations for *M. acuminata* (t=251, df=186.17, p=0.026), and *M. balbisiana* (y=4.720, df=84.176, p<0.001), but not for *M. maclayi* (t=1.216, df=137.97, p=0.226).

### Species level differences

Genetic profiles were characteristically distinct according to species. Loci polymorphism for *M. acuminata* was on average 3.84, for *M. balbisiana* 6.18 and for *M. maclayi* 18.64 (Table S3). In all cases H_o_ was less than expected heterozygosity (Nei’s gene diversity, H_exp_) apart from *M. balbisiana* populations where they were approximately equal. The disparity was especially evident for *M. acuminata* population and seeds, and *M. balbisiana* seeds. Evenness (E_5_) was high (>0.9) across all sampling groupings but less so for *M. acuminata* seeds (E_5_=0.684). *Musa acuminata* seeds and populations had the lowest diversity in all sampling groups. Fewer multilocus genotypes were observed in *M. acuminata* populations and, to a greater extent, seeds, compared to the number of actual samples - suggesting clonality or self-fertilization of homozygous mother plants; this was also observed in *M. balbisiana* seeds but to a lesser extent. Heterozygosity was very low in *M. acuminata* and *M. balbisiana,* but high for *M. maclayi.* There was no evidence of null allele excess, large allele drop out and error due to stuttering in *M. balbisiana* or *M. maclayi*. Six out of the 19 loci for *M. maclayi* showed potential null allele excess, possibly inflating homozygosity beyond predicted values, however, homozygosity was high across all loci (Table S4).

### Representation of populations in seeds at the local level

Allelic richness of local seeds as a proportion of the local populations from where they were collected was 56±20%, 76±42% and 78±18% (mean and standard deviation, *M. acuminata, M. balbisiana* and *M. maclayi* respectively, Table 3).

**Table 3.**
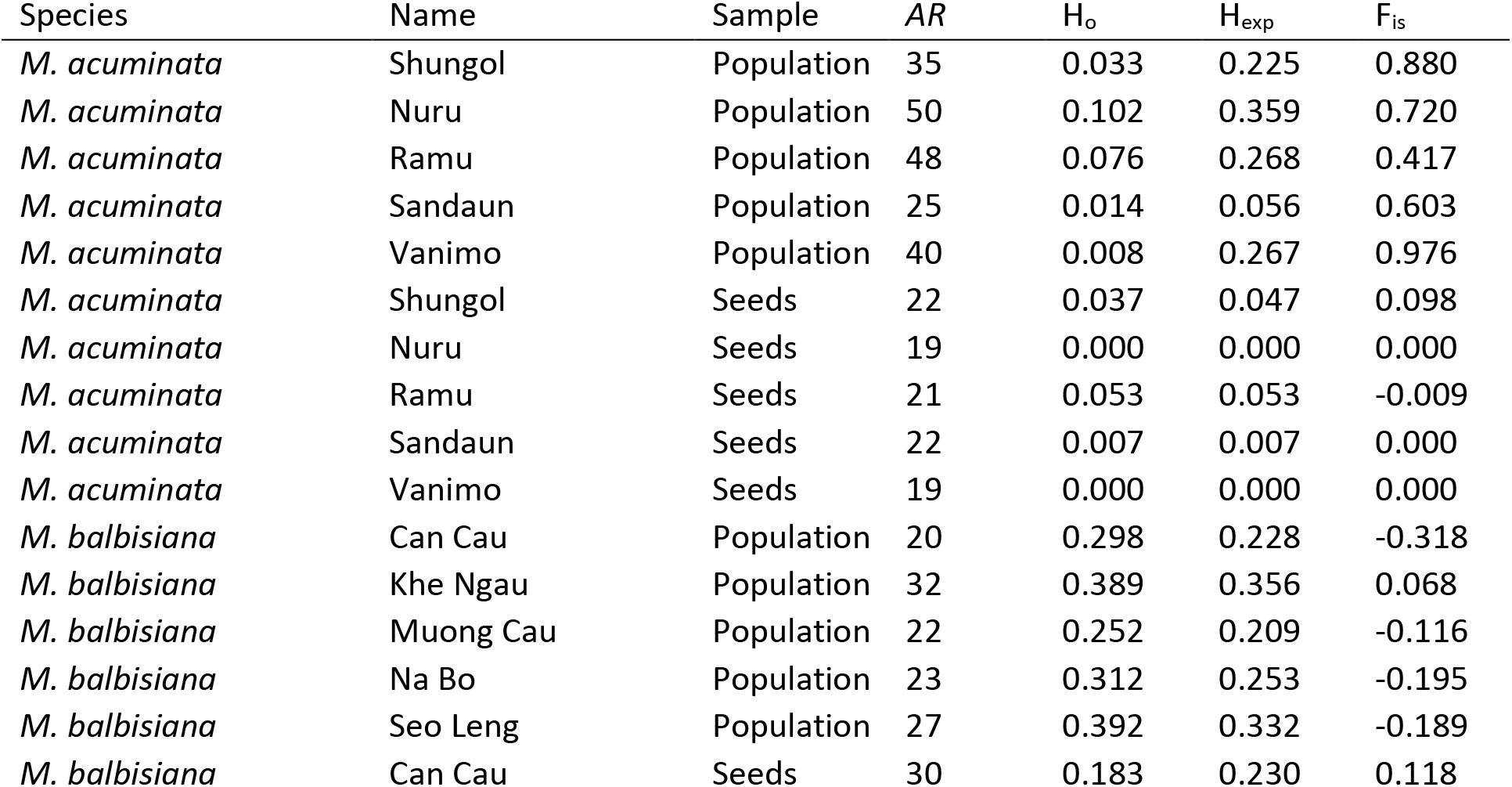
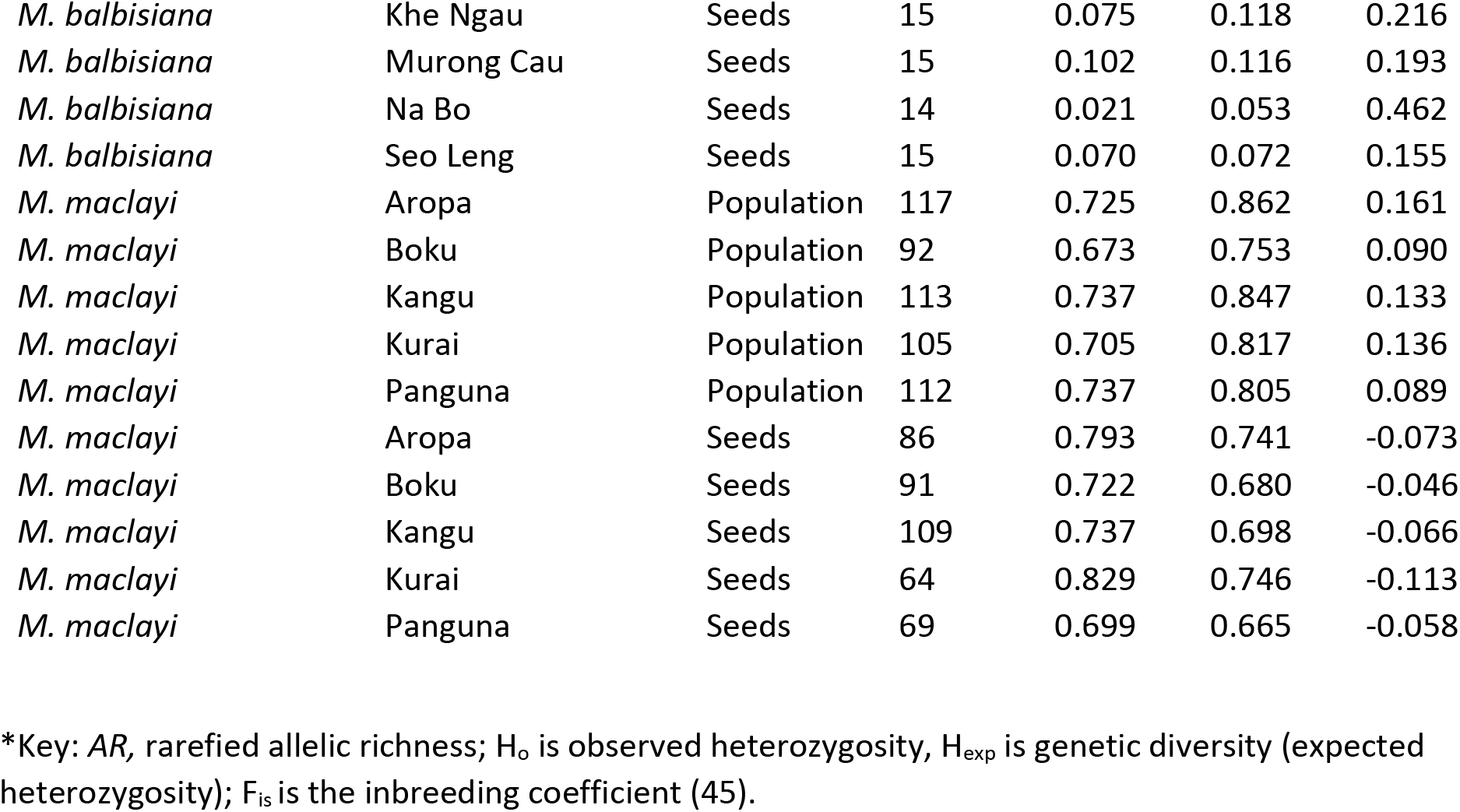
Diversity indexes of populations and seeds pooled at the local level*.

*M. acuminata* seed collections had very low Ho including two bunches where H_o_ was zero (Table 3). H_exp_ varied considerably in local populations of *M. acuminata*. Nuru was the most diverse (H_exp_=0.36), notably seeds from this population were completely homozygous (H_o_=0). Sandaun was the least diverse *M. acuminata* population (H_exp_=0.06). Inbreeding coefficient (F_is_) was high for all *M. acuminata* populations, Vanimo being the most inbred (F_is_=0.97), its seeds were also completely homozygous. The Ramu population was the least inbred (F_is_=0.41), and had seeds with the highest heterozygosity (H_o_=0.05). *Musa balbisiana* populations are also characterized by a low degree of diversity, yet inbreeding coefficients were much lower compared to *M. acuminata*. By contrast, populations of *M. maclayi* were characterized by a high level of heterozygosity and genetic diversity and low inbreeding coefficients. Populations of *M. balbisiana* and seeds of *M. maclayi* had negative Fis meaning an excess of heterozygotes.

### Targeting local seed collections

Allelic richness of both the regional population and bunches was estimated by bootstrap resampling to estimate the total pool of alleles including unseen alleles. To capture 70% of alleles estimated to be present in the region, at least four bunches need to be collected for *M. balbisiana* (Figure2b); for 90% of alleles, at least five bunches are necessary. For *M. maclayi* three bunches are needed to sample 70% of regional *AR*, and four bunches for 90%, despite the much higher *AR*. Allelic sampling for *M. acuminata* had a different profile. It was not possible to collect even 70% of regional *AR*, and for each bunch collected there was minimal gain in allelic capture. If many more bunches were sampled it may be possible to capture up-to 70% regional *AR*, but this would probably require >15 bunches, based on extrapolation (Figure 2b).

Alleles in seeds of *M. acuminata* are largely shared by all bunches and by the regional population (Figure 3). Each cumulative bunch adds only a few alleles. Populations with minimal overlap, and therefore maximum coverage, include combinations of Ramu and Vanimo, and Ramu and Nuru. *Musa balbisiana* displayed a similar overlapping pattern to *M. acuminata* in that many of the alleles in seeds were shared by bunches. Seeds from the Can Cau population had the least shared alleles, with very little gain each time a bunch was added. Bunches from *M. maclayi* had less overlap, suggesting genetic structure and isolation by distance. No alleles in bunches were shared by all bunches, a large proportion of regional alleles was covered by bunch ellipses.

**Figure 3.**
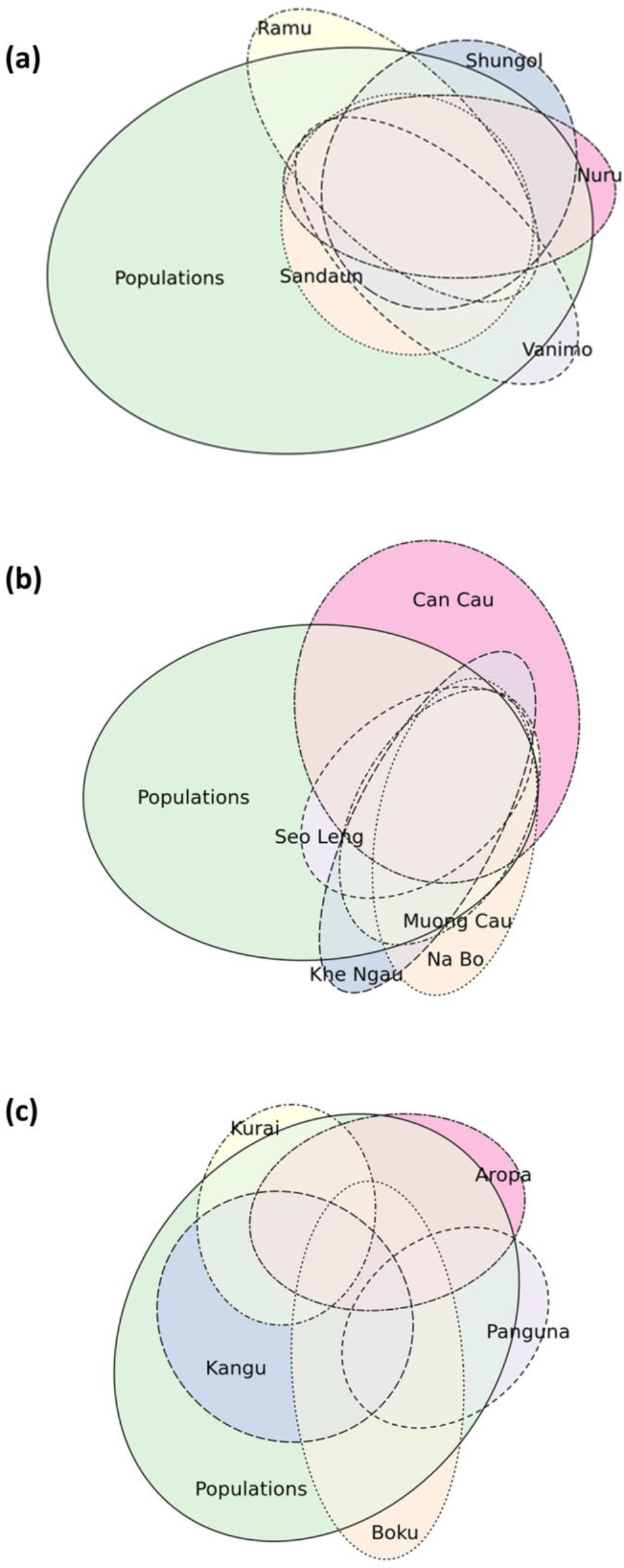
Euler plots of allele grouping in local seeds and regional populations, size and overlap of ellipses is relative to the number of alleles and the amount they share with other groupings: (a) *M. acuminata* (b) *M. balbisiana* (c) *M. maclayi*

Genetic distance (96) was calculated pairwise between all local populations and local seeds (Figure 4). For *M. acuminata* genetic distance was low between all samples. Populations and seeds from Vanimo were most distant from other samples. Seeds clustered with their respective populations for Vanimo and Sandaun, but not for other populations of *M. acuminata*. For *M. balbisiana,* seeds from Na Bo and Seo Leng were most distant. Notably the outlier population of Khe Ngau was not more distant from other populations and seeds. Several populations and seed pairs of *M. maclayi* clustered together. There were two broad clusters with samples from the North West (Boku and Panguna) having greater distance from the other three populations. Isolation by distance was evident in *M. maclayi* populations and seeds (Mantel test, 999 permutations; populations, p=0.03; seeds p=0.013), but not the other species.

**Figure 4.**
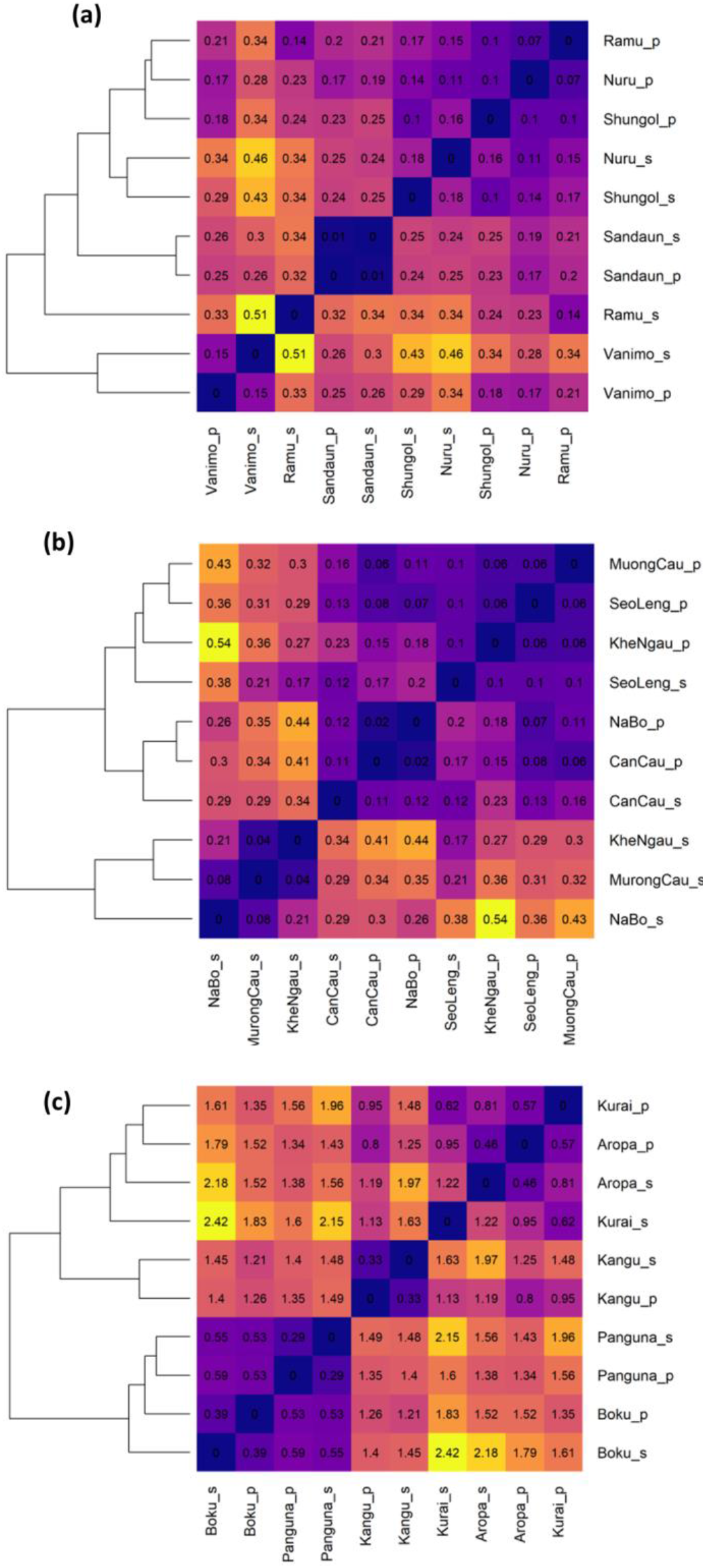
Pairwise genetic distance (Nei, 1972) of populations and seeds, and clustered dendrograms using hierarchical clustering: (a) *M. acuminata* (b) *M. balbisiana* (c) *M. maclayi*

### Selecting seeds from the same local collection

Accumulation of *AR* of seeds per bunch was estimated (Figure 2c). For *M. acuminata,* a single seed contained over 90% of the *AR* of the whole bunch. The allele accumulation curve is virtually flat, seeds are therefore more-or-less genetically identical. For *M. balbisiana* over 70% of alleles are found in only two seeds, and 90% of alleles in 10 seeds. For *M. maclayi,* 70% of the estimated total alleles in the bunch are captured by 16 seeds. To achieve 90% of the total the accumulation trend line must be extrapolated considerably beyond the data to around 35-50 seeds.

## Discussion

### Genetic capture in seed collections compared to their source populations, for three focal species

Allelic richness of seeds as a proportion of populations met the conservation target of 70% from the Global Strategy for Plant Conservation (2) for two out of the three focal species (*M. balbisiana* 81% and *M. maclayi* 93%). *Musa acuminata* only achieved 51% proportional allelic richness. In all cases several seed collections were required from different local populations to maximize genetic capture. The number of collections necessary to achieve the 70% target was species dependent. For *M. acuminata* it was >15 local seed collections, for *M. balbisiana* four and *M. maclayi* three.

All common alleles of populations were included in seed collections, but the level of representation was lower for rare alleles, with some rare alleles missing from seed collections (Figure 2a). Brown and Marshall’s sampling strategy (15), used by about two thirds of leading seed conservation institutions (13), advocate sampling 30 individuals for out-crossing and 59 for selfing species, with the aim of having a 95% chance of capturing alleles with frequency <0.05. The results of the present study show that collecting from a much lower number of mother plants (a total of five) resulted in relatively high genetic capture, even including alleles rarer than the threshold set by Brown and Marshall. Furthermore, against the key success criteria proposed by Brown and Marshall (15) - capturing locally common alleles (because globally common alleles are easily collected in any sample and globally and locally rare alleles are ultimately limited by the sample size) - our results show that seed collections of *M. maclayi* and, to a lesser extent, *M. acuminata* and *M. balbisiana,* were successful (Figures 3 and 4).

A high degree of homozygosity and low level of diversity, apparent in populations and seed collections of *Musa acuminata* subsp. *banksii* in our study, as well as that of Christelova et al. (46), corroborate a typical genetic signal that is associated with self-fertilization (47). Unlike cultivated and most wild banana species (including other *M. acuminata* species), seed bearing *M. acuminata* subsp. *banksii* are characterized by self-compatible hermaphroditic flowers, particularly in the upper hands on the inflorescence (32, 38), these likely self-pollinate prior to flower bract opening (48). Similar floral morphology, and therefore probably self-pollination, is also observed in *M. acuminata* var. *chinensis* (49), *M. boman* (30), *M. jackeyi* (interestingly closely related to *M. maclayi*) (32), *M. ingens* (30), *M. rubinea* (50), *M. schizocarpa* (30), *M. yunnanensis* (49) and *M. zainful* (51). Seed collections derived by self-fertilization are naturally more representative of the mother plant than the population. Therefore, to capture the genetic diversity in populations of self-pollinating *Musaceae* species, many mother plants must be sampled.

By contrast, populations and seeds of *M. maclayi* were characterized by higher levels of heterozygosity and diversity, consistent with cross-fertilization (52). Male and female flowers of *M. maclayi* are temporarily and physically isolated on the same inflorescence. Female flowers are produced first, followed by male flowers, as the peduncle grows (30). Genetic capture in *M. maclayi* seeds therefore represent both the mother plant and pollen donors within the population. As a result, less bunches need to be collected to represent the population compared to *M. acuminata*. The level of diversity evident in the small number of populations of the present study was somewhat surprising, because of the narrow distribution and relatively recent diversification and dispersal of the former Australimusa group to which *M. maclayi* belongs (53). This demonstrates the strong effect of mating system on genetic diversity in populations and seeds.

Populations of *M. balbisiana,* in our results, had low heterozygosity and diversity, this is in keeping with several previous studies (24, 35, 46, 54, 55). Moreover, the heterozygosity of seed batches was much lower than within populations. Our results were similar to those found by Bawin et al. (56) for *M. balbisiana* seeds collected from *ex situ* field collections or feral populations, but our seeds were less diverse than those collected from native populations in Yunnan (China). Even though *M. balbisiana* basal flowers are functionally female (30) and do not produce seeds when pollinators are excluded (36), flowers may effectively be selfed from a different flower of the same genotype on the same mat or from vegetatively reproduced neighbouring plants (57).

The low diversity in populations of *M. balbisiana,* in our results, may be caused by a genetic bottleneck and/or founder effect. This hypothesis was also proposed by Ge *et al.* (35) and Shepherd (58) and is in keeping with *Musa* ecology: being early successional or disturbance-adapted (59, 60). Additionally, the intensive deforestation and reforestation that has occurred in Viet Nam over the past 50 years (61) may also be causal. Indeed, according to a recent study (62), the ecological traits of *M. balbisiana* makes them particularly vulnerable to genetic erosion from anthropogenic disturbance. Furthermore, as *M. balbisiana* has many uses by local communities (63), plants are often planted or encouraged in vacant land. Finally, seed collections may indeed result from introgression from neighbouring cultivated bananas. These possibilities illustrate some of the challenges associated with conservation of CWRs by seed.

Variation in genetic capture of different species of the *Musa* genus demonstrates the profound effect of mating system on genetic capture in seed collection. Taxonomically relatedness, therefore, is not a good proxy for a sampling strategy (64). In support of our results, a recent study by Hoban *et al*. (65) found species in the same genus required on average 50% more individuals to reach desired levels of capture compared to others. Furthermore, depending on mating system, dispersal distance, life cycle and the sampling strategy employed - up to 5 times as many individuals may need to be sampled for the same level of genetic capture (14).

### Guidance about how to maximize genetic capture for future seed collections

To maximize genetic capture in *Musa* seed collections, firstly, we recommend that species mating systems should considered to inform sampling strategies. Our results are therefore in support of Brown and Marshall’s sampling strategy discussed above (15).

For self-pollenated *Musa* species, as many mother plants should be sampled from as possible. For species with wide distributions, populations should be spatially dispersed; however this is less important than increasing the number of plants collected from. Collecting seeds from many individuals of adequate quality for long term storage is highly challenging; it is not straightforward to find mature seeds in the forest suitable for storage (66, 67). It would certainly not be possible to collect from the 59 individuals proposed by Brown and Marshall (15), or even the 15 proposed here, in one collecting trip. As bananas fruit throughout the year, seed collections may therefore require repeated temporal sampling.

To target collections of fully out-crossing species, fewer collections are required to represent regional alleles. We recommend collections should be focussed on increasing the number of local populations collected from rather than the number of mother plants in a population. Local populations should be spatially dispersed to maximize genetic capture. This will also allow for locally distributed alleles to be captured (15). The amount of both rare and locally distributed alleles therefore depends on resources for collection, but there are diminishing returns associated with such effort.

For all species, but especially for out-crossing species, it is also important to target collections that are far from agriculture and human interference. Large and well established populations should be prioritised (62). This will likely maximize genetic diversity in source populations (68), and avoid unwanted introgression from cultivated forms (69).

### Direction for seed distribution on how to provide representative seed samples

To ensure enough seeds are conserved, self-pollenated species only require one or two seeds from a bunch to be part of a core collection. There is also very little point in using many samples of self-pollenated seeds in experiments. This contrasts with fully out-crossed seeds, where more seeds should be conserved in core collections per bunch or used as samples in experiments. For *M. maclayi* 16 seeds represent 70% of alleles, and 35-50 seeds represent 90% of alleles. Even so, these numbers of seeds are easily achieved, for most *Musa* species at least, where a bunch can contain hundreds to thousands of seeds. However, for some species we have collected (e.g. *M. ingens*), only a few seeds were found in a bunch, perhaps due to inadequate pollination. Additionally, these findings mean that despite low levels of survival in storage of some collections (59, 66), population genetic diversity can be protected in a few seeds.

### Limitations

The present study was constrained in that only one mother plant was used per local population, and only 5 per region. It was therefore not possible to test the effect of additional local seed collections on genetic capture. This was because accessing bunches at the right level of maturity for germination and storage is one of the key challenges for seed conservation of banana CWRs (66); often mature bunches are not to be found in a forest population. Furthermore, in the present study we compared genetic capture in seed collections at the regional level. This does not account for the full level of diversity across species distributions which may be much wider than that sampled here, particularly in the case of *M. balbisiana* (Table 1). Further research should be done to assesses isolation by distance of source populations and seed genetic capture to optimise sampling strategies that use species distributions across ecozones as sampling strategies (e.g. 70). Additionally, sampling did not consider temporal effects in sampling, such as collecting from the same populations at different time points, this may prove important, at least for cross-fertilized species.

It is also important to emphasize that whilst broad comparisons between species are of interest, direct comparison between species from our results should be cautioned because different taxon-specific microsatellite markers were employed. Observed allelic variation may indeed be resultant of specific markers used, rather than actual differences, meaningful at species level. However, as we used suites of 16-19 markers per species in the present study this effect is minimised, despite this, any comparative interpretation should be taken with caution. Importantly, direct comparisons between species was not our primary purpose, rather, our aims were to assess genetic capture in seed collections compared to their source populations for three focal species.

## Conclusions

Seed banks are efficient ways of conserving genetic diversity in wild populations and make it available for future use in breeding programmes or conservation. However, because very little is known about both population and seed genetic diversity the representativeness and therefore the value and use of seed collections is limited. We have demonstrated the measurement of genetic capture in seed collections of three of the most important wild relatives of the most important fruit crop in the world. We have shown how targeted seed sampling should be species specific and genetically informed; notably, species mating systems and evolutionary history (whether natural or anthropogenic) have a profound effect on the level of genetic diversity in seed collections. The results of the present study may be applied across sampling strategies of other wild species, in that species mating systems should be a primary consideration to maximize genetic representation in seed collections.

## Methods

### Focal species

We focused on three wild *Musa* species: *M. acuminata* subsp. *banksii* (F.Muell.) N.W. Simmonds*, M. balbisiana* Colla and *M. maclayi s.l.* F.Muell. (hereafter termed *M. acuminata, M. balbisiana* and *M. maclayi*) (see Table 1). In this study *M. maclayi s.l.* includes closely related *M. bukensis* and *M. maclayi* subsp. *maclayi* taxa that occur on the island of Bougainville. Based on the description of both taxa and personal observations there is evidence of introgression between the two taxa on the island, and it is unclear whether they are two different, or one single, species (30).

### Study region and populations

Natural populations of focal species in their respective native ranges were sampled during several collecting missions that took place between 2016 and 2019 (Figure 1; Table 1). Collection of *M. acuminata* was carried out in Papua New Guinea (PNG) in June 2017 and May 2019 (66, 71). *M. balbisiana* was collected in Viet Nam during November 2018 and April 2019. *Musa maclayi* was collected on the island of Bougainville (PNG) in October 2016 (72, 73).

### Plant material

Leaf and seed samples were collected from wild natural populations. All seeds, leaves and data were collected and transferred according to local legislation and supplied for non-commercial use and research under the Standard Material Transfer Agreement in accordance with the International Treaty on Plant Genetic Resources for Food and Agriculture. None of the species included in the present study are CITES listed. Formal field identification was carried out by Steven B. Janssens (Meise Botanic Garden, Belgium). Leaf samples were collected randomly from 5 local populations per species (Figure 1, Table S1). From each population, leaves from 15 plants on average were sampled and further used in this study. Dried leaf samples were taken to the laboratory following the field mission for DNA extraction. A single seed containing infructescence (hereafter termed ‘bunch’) was also collected from each population. Groups of fruits (hands) from the former clusters of flowers subtended by one bract, were separated and processed separately after shipping to Meise Botanic garden as described by Kallow et al. (66). Bunches collected in Viet Nam were not separated by hand and were processed in a similar way in the laboratory of Plant Resource Center (Ha Noi, Viet Nam). In both cases, seeds were stored at 15% relative humidity and −20 °C prior to germination and DNA extraction.

To overcome barriers associated with low and unpredictable *in vivo* germination, seeds were germinated by embryo rescue as described by Kallow et al. (66). Seeds were selected randomly from 2-3 hands per bunch, or, for 3 bunches of *M. acuminata* and all bunches of *M. balbisiana,* from pooled seeds from the whole bunch. Due to low seed numbers and viability of *M. balbisiana* accessions DNA was extracted directly from their embryos. An average of 16 seeds per bunch were used in this study.

For each population, exact coordinates were recorded with a Garmin GPS device. Detailed taxonomic field notes and notes on geography and ecology were recorded for each sample. Photographs of mother plants (the plant from which the bunch was taken) and of bunches were taken. Seed samples from PNG and Bougainville were accessioned into the Meise Botanic Garden seed bank (Meise, Belgium). Seeds from Viet Nam were accessioned into the seed bank of Plant Resources Center (Ha Noi, Vietnam).

### Microsatellite PCR

We isolated DNA using a method adapted from Doyle and Doyle (74) and then sequenced samples using a suite of taxon specific polymorphic microsatellite markers arranged in multiplexes (Table S2). For *M. acuminata* we developed mutiplexes from previous studies (35, 75–80). A total of 86 primer pairs were tested for amplification individually and then arranged in a total of 15 multiplexes using Multiplex Manager (81) and Multiple Primer Analyzer (82) with 12 *M. acuminata* samples. From this, 20 markers arranged in four multiplexes were selected. For *M. balbisiana,* we used the multiplex arrangement of Bawin et al. (56). These included 18 SSR markers organized into four multiplexes. For *M. maclayi,* a total of 16 specific SSR markers were newly developed and optimized by Genoscreen (Lille) and arranged in four multiplexes. We used an M13 labelling protocol (83) to arrange multiplexes. We used the Type-it Microsatellite PCR Kit (Qiagen, Venlo, the Netherlands) to amplify microsatellite regions. We then sequenced the resultant PCR product on an ABI 3730 sequencer (Applied Biosystems, Foster City, California, US). See Supplementary Methods for detailed methodology.

### Data analysis

#### Fragment length analysis and quality check

We analyzed microsatellite fragment lengths using Geneious v 8.1.9 software. Loci and samples with more than 25% missing data were excluded from the analysis to allow for missing-ness to be similar for seeds and populations. This resulted in excluding from the data 1 locus used for *M. acuminata* data, 8 for *M. balbisiana* and 2 for *M. maclayi* (Table S3). Several loci were missing from *M. balbisiana* presumably because of low DNA concentrations resultant of extraction from embryos rather than leaves. Resultant missing data was 3.9% for *M. acuminata,* 8.1% for *M.balbisiana* and 4.7% for *M. maclayi.* We then assessed allele data for null allele excess, large allele drop out and error due to stuttering using the Microchecker software (84).

### Genetic assessment

Genetic assessment was carried out at two levels: the regional level whereby samples were pooled by either all local populations or all seeds per species; and the local level whereby samples were not pooled but kept separate from each local population and each bunch per species. At the regional level we calculated several indices to represent genetic diversity of populations and seeds. All computations were carried out in the R environment (85). As a broad estimate of the amount of genetic material present, we determined *AR* rarefied to equal sample size (86), using the *pegas* package (87). We counted *PA*, present in populations and not seeds and *vice versa* using the *poppr* package (88); and, in order to assess the rarity of alleles in samples, assessed the relative frequency of alleles, computed in the *adegenet* package (89). To represent the genotypic diversity of samples and to assess inbreeding we calculated H_exp_ (44). We also measured the Ho to assess population genotypic diversity and inbreeding. The number of MLG was computed, as an indicator of clonality. Several commonly used diversity indices were also calculated using the *poppr* package (88): H’, λ and E_5_. At the local level, we repeated calculations of *AR*, H_o_,H_exp_ and additionally calculated Fis (45) in the *hierfstat* package (90). Indices were compared using two-sample t tests.

### Cumulative proportional allelic richness

We assessed how many bunches required to capture 70% (based on Target 9 of the Global Strategy of Plant Conservation) (2) and 90% (an arbitrary but sometimes used threshold) of alleles in the region per species. We did this by firstly calculating the regional *AR* and then *AR* of bunches which was then added cumulatively by taking the mean and standard deviation of each cumulative step. To also assess how many local populations (rather than seeds) should be sampled to represent the regional population, the same analysis was completed by cumulatively adding *AR* of local populations. Estimates were normalised as a proportion of extrapolated regional *AR* that was estimated using bootstrap resampling (91). We also used this method to estimate *AR* of local populations and bunches.

A similar approach was employed to estimate proportional cumulative *AR* of seeds per bunch. For this, total *AR* of bunches was extrapolated (92), and then seeds were added cumulatively as described. Computations were made in the *vegan* package (93). Trend lines were plotted using the loess method in ggplot2 (94).

### Allele groupings and genetic structure

We made allele groupings for local seeds and regional populations (http://bioinformatics.psb.ugent.be/webtools/Venn) and plotted them as Euler diagrams using the *eulerr* package (95). We calculated pairwise genetic distance of local populations and seeds(96), and produced a heat map with dendrogram using complete linkage hierarchical clustering. We then assessed isolation by distance by comparing Euclidean distances of coordinates and population matrices of seeds and populations (separately) using the Mantel test.

## Declarations

### Ethics approval and consent to participate

All seeds, leaves and data were collected and transferred according to local legislation and supplied for non-commercial use and research under the Standard Material Transfer Agreement in accordance with the International Treaty on Plant Genetic Resources for Food and Agriculture. None of the species included in the present study are CITES listed.

### Consent for publication

Not applicable

## Abbreviations

*AR*: allelic richness
CITES: Convention on International Trade in Endangered Species of Wild Fauna and Flora (1973)
CWRs: Crop wild relatives
df: degrees of freedom
DNA: DeoxyriboNucleic Acid
E5: evenness index
Fis: inbreeding coefficient
GPS: global positioning system
H’: Shannon:Weiner diversity index
H_exp_: Nei’s gene diversity (expected heterozygosity)
H_o_: observed heterozygosity
MLG: multilocus genotypes
*Musa acuminata*: Musa acuminata subsp. banksii
*Musa maclayi*: Musa maclayi s.l.
N: number of samples
*PA*: private alleles
PCR: Polymerase chain reaction
PNG: Papua New Guinea
Species: species and infraspecifics
*Λ*: Simpson’s index

## Availability of data and materials

The datasets used and/or analysed during the current study are available from the corresponding author on reasonable request.

## Authors’ contributions

SK: Conceptualization, data curation, formal analysis, investigation, methodology, software, visualization, writing – original draft, writing review & editing. BP: conceptualization, funding acquisition, project administration, resources, supervision, writing – review & editing. ToDV: data curation, funding acquisition, resources. TuDV: data curation, resources. JP: resources. AM: data curation, resources, validation, writing – review & editing. RS: validation, writing – review & editing. SBJ: conceptualization, funding acquisition, methodology, project administration, resources, supervision, validation, writing – review & editing.

## Acknowledgements

We gratefully acknowledge those who carried out the seed and leaf collections. In PNG and Bougainville: J. Pilon, I. Nabo, B. Pitalai, G. Savi, P. Daur, E. Yabu, S. Itau, J.Lapiu, T. Kunou, S. Kambase, J. Guaf, N. Sinoksor, S. Carpentier, J. Sardos, G. Sachter-Smith and D. Eyland. In Viet Nam: Le Thi Loan, Ngo Duc The. Thank you also to T. Vanderstraeten, K. Longin, N. Fanega Sleziak and H. Krohn for carrying out embryo rescue and DNA isolation. We are grateful for technical laboratory work and helpful advice from W. Baert, P. Asselman, Y. Bawin, A. Heylen and S. Vanden Abeele at Meise Botanic Gardens. We also gratefully acknowledge S. Hoban and R. Gargiulo for advice on the analysis and manuscript and J. Sardos for the basis of Table 1.

